# Improved FRAP measurements on biofilms

**DOI:** 10.1101/707950

**Authors:** J. Hauth, J. Chodorski, A. Wirsen, R. Ulber

## Abstract

We expand the standard FRAP model introduced by Axelrod et al. in 1976. Our goal is to capture some common artifacts observed in the fluorescence measurements obtained with a confocal laser scanning microscope (CLSM) in biofilms: 1) linear drift, 2) exponential decrease (due to bleaching during the measurements), 3) stochastic Gaussian noise, and 4) uncertainty in the exact time point of the onset of fluorescence recovery. In order to fit the resulting stochastic model to data from FRAP measurements and to estimate all unknown model parameters, we apply a suitably adapted Metropolis-Hastings algorithm. In this way, a more accurate estimation of the diffusion coefficient of the fluorophore is achieved. The method was tested on data obtained from FRAP measurements on a cultivated biofilm.

**STATEMENT OF SIGNIFICANCE:** Diffusion and mass transport in biofilms is presumed to play an important role in the resistance against antimicrobial agents and the secretion thereof. FRAP measurements give insight into these transport processes. In this article, the authors expand the standard FRAP model by a blackbox part which addresses some artifacts commonly observed in the measurements. This is done in order to improve the estimations of diffusion coefficients considerably, aiming at a more comprehensive description of diffusion processes inside the biofilm. We expect that the methods are transferable to FRAP measurements on materials other than biofilms.

## INTRODUCTION

Diffusion and mass transport in biofilms has been studied for decades, yet has still not been understood completely. It is presumed to play an important role in the resistance against antimicrobial agents and the secretion thereof (1). However, there are different ways of mass transport into and inside a biofilm. First, the studies of mass transport inside the biofilm from the bulk medium are more numerous than the studies investigating transport inside the biofilm (2–7). Second, there is a huge emphasis on oxygen transport in the biofilm (8–11). But how does diffusion inside of the biofilm work? How are nutrients transported from one part of the biofilm to another, if any? How are products excreted? One way to look inside a biofilm is by usage of the confocal laser scanning microscope (CLSM) and the fluorescence recovery after photobleaching (FRAP) technique. An important challenge here is the visualization of the transported compounds. Since most of the substrates and products from microbial biofilms (e.g. glucose and lactic acid) don’t have any viably detectable fluorescence, other substances have to be used in order to be able to perform the FRAP experiments. In 1974, Peters et al. (12) started using fluorescein isothiocyanate (FITC) as a marker for diffusion measurements. Since then, it has been widely used in many different studies (13–17). The first model for describing mobility of compounds by FRAP was developed by Axelrod et al. in 1976 (18). Since then, several improvements were created for this basic model, such as simplified computations or using the ratio between bulk fluid diffusion and biofilm diffusion (19–21), as well as the technique itself (22, 23). In this study, the authors build upon these models and expand them in order to improve the estimations of diffusion coefficients, aiming at a more comprehensive description of diffusion processes inside the biofilm.

The principle behind FRAP is based on the idea that after the induction of an irreversible bleaching of a fluorophore with a strong laser pulse, the subsequent observation of the recovery of the fluorescence in the bleached region reveals insight into the processes governing transport and diffusion of this fluorophore. Since the fluorescence recovery can only originate from fluorescing material which diffuses into the bleached region from its surroundings, the development of the recovery and the corresponding increase in fluorescence intensity allow the model-based estimation of e.g. the diffusion coefficient of the fluorophore. Finally, this in turn gives insight into the transport processes inside the system under observation (23, 24).

An important step in the analysis of each FRAP experiment is the evaluation of the measurement data which has to be based on a mathematical model. In the past, these models were kept very simple in order to guarantee that their solutions could be provided by simple explicit formulas (16–18, 21, 25). Nevertheless, the restrictive model assumptions necessary for the explicit solvability of these models made by Axelrod or others building on this model (19, 22) are often problematic or even not valid for the real systems under consideration. This is especially true in the case of biofilms. The simplifying assumptions usually made are e.g. that the bleaching during the measurement (in the post-bleaching phase) is neglected, that the probe is homogenous at the region of interest, that the processes in the third dimension are neglected, and that the laser profile has a very simple geometry (Gauss profile or circle) (26). Apart from these properties which are inherent in the real system and well understood, there may be additional artifacts in the measurements like gradients in the fluorescence intensity or steady changes of it even if no bleaching is done. In order that the mathematical analysis achieves better results, it is therefore necessary to deal with more complex models. The downside is that these models do not yield solutions given by simple explicit formulas.

In this article, we expand the standard FRAP model by a blackbox part which addresses some artifacts commonly observed in the measurements but which violate the assumptions made for the standard FRAP model. In detail, the model will capture the following disturbances of the fluorescence measurements: 1) a linear drift, 2) an exponential decrease (due to bleaching during the measurements), 3) stochastic Gaussian noise, and 4) uncertainty in the exact time point of the onset of fluorescence recovery. We apply a Markov Chain Monte Carlo algorithm in order to fit the expanded model to data obtained from FRAP measurements and to estimate all unknown model parameters, including the diffusion coefficient. We expect that the methods are transferable to FRAP measurements on materials other than biofilms.

## MATERIALS AND METHODS

### Biofilm cultivation and microscopy

#### Reactor construction

Biofilms were cultivated in a flow cell. The flow cell (FC) used in this study was a modified version of a custom-made flow cell described by (27). For application of fluorescent dyes, an inlet pointing at the sample well was added (Fig. 1) to ensure homogenous distribution of the applied dye. The height of the flow channel was decreased to 1.6 mm, to a total reactor volume of 1.5 mL, allowing full coverage of the biofilm volume with the given objective. CFD (Computational Fluid Dynamics) simulation (data not shown) proved the flow to be laminar.

**Figure 1:**
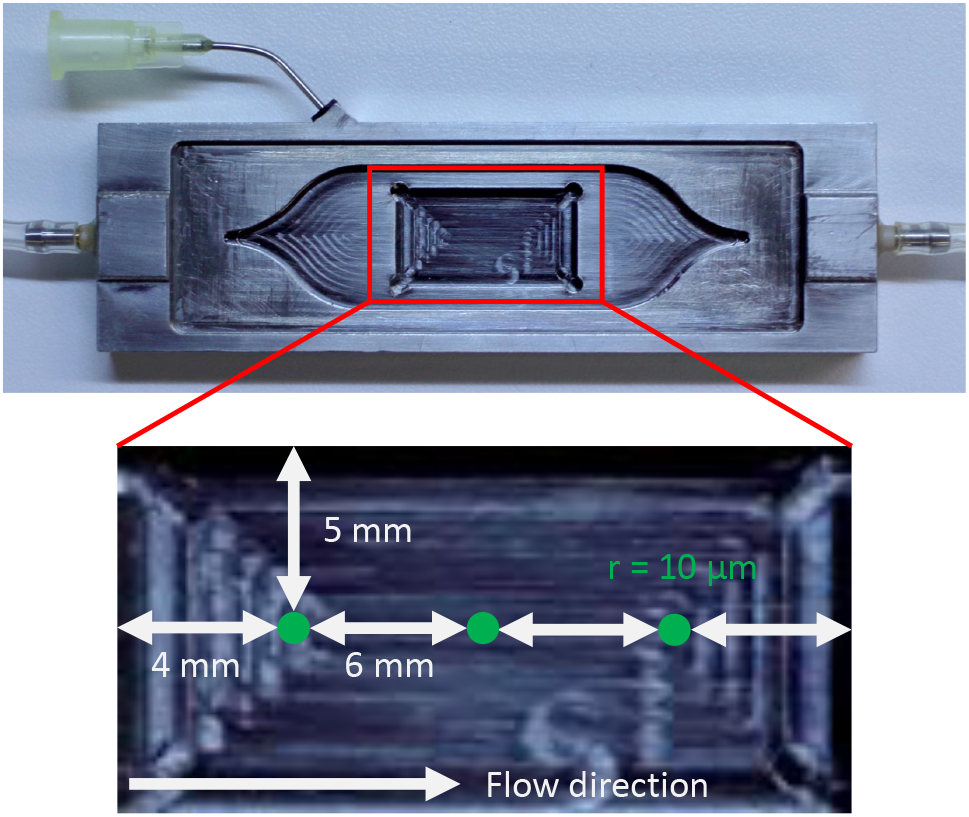
**Top:** FTC with dye inlet and sample well. **Bottom:** Measurement positions on a 10 x 20 mm metallic surface fit into the well of the same size in the FC. Positions are evenly spaced 5 mm from the long edge, and 4 mm (upstream/downstream) or 10 mm (center) from the short edge, respectively. ROI is 20 *μ*m in diameter.

#### Bacterial cultivation

For this study, biofilms of the lactic acid producing organism *Lactococcus lactis subsp. lactis* (DSMZ 20729) were generated. All cultivations were carried out at 30°C in MRS (de Man, Rogosa, Sharpe) medium (28) and were split into three phases – A) Pre-culture, 100 mL MRS, pH 7, 20 g/L glucose, in a bottle shaken at 130 rpm to an OD_600_ (Optical Density at 600 nm) of 4, which was then used to inoculate the batch culture. B) Batch culture, with the medium being recycled through the FC in order to allow for bacterial attachment and initial biofilm formation: 100 mL bottle connected to FC, pH 7, 20 g/L glucose, pumped at 5 mL/min for 15 min with 60 min break between each interval (Ismatec IPC peristaltic pump, IDEX Health & Science, Wertheim, Germany) for 16h. C) Continuous culture, 250mL MRS, pH 5.5, 10g/L glucose, 0.25 mL/min flow for up to 12 h, during which all measurements were carried out.

#### FRAP measurements

FRAP measurements were carried out on a Leica SP5 II confocal laser scanning microscope (Leica Microsystems, Wetzlar, Germany) with an active beam expander module. In the first eight hours of biofilm growth in continuous mode, pictures of the surface were taken at 0h and 3 h. Starting with the 5 h timepoint, 100 *μ*L fluorescent dye was added. Starting at the 7 h timepoint, dye was applied every hour until 12 h. The dyes used were uranin (VWR International GmbH, Darmstadt, Germany) at a concentration of 0.75 mg/mL to an end concentration in the reactor of 0.05 mg/mL and FITC Dextran 4kDa (FD4) (TdB Consultancy AB, Uppsala, Sweden) at a concentration of 225 mg/mL to an end concentration in the reactor of 15 mg/mL. Both dyes are not known to have any biological effect on or reaction with prokaryotes (Uranin: see (29), FD4: widely used, no reports of adverse effects). Pumping was continued for two minutes after application to allow for the dye to spread across the whole reactor volume and was then shut off in order to not disturb measurement. FRAP time series were taken with a 63×0.9 water immersion objective in the biofilm at a set distance from the substrate, as well as in the bulk fluid above the biofilm. The measurement parameters were as follows: 256 px image size, 1400 Hz scanning speed (bidirectional), 6x zoom, 20 *μ*m circular bleach radius, 75 % laser power (Argon laser), 5 % AOTF (acusto-optical filter) (scan)/100 % AOTF (bleach), pinhole 2 AU (Airy Units). Each measurement was separated into a pre-bleach, bleach and post-bleach phase. Pre-bleach 15 frames, bleach 5 frames, post-bleach 60 frames. Frame time was 0.195 s. 8-bit TIFF (Tagged Image File Format) was used as data format. Three positions for measurement were chosen on the 10 x 20 mm sample, as shown in Fig. 1.

### Mathematical Modeling of FRAP: Review

The FRAP model was introduced and solved for circular and Gaussian laser profiles by Axelrod and his coworkers (18). Soumpasis (19) corrected a typo in that paper (in the main formula (14), the Bessel function *I*_2_ should be *I*_1_) and developed a simplified and numerically more stable formula for circular profiles. Before presenting our modifications, we restate the model developed in (18) and (19).

The fundamental model describes the time development of the spatially distributed fluorophore concentration in reaction to a certain laser intensity profile. A two-fold reaction takes place: fluorescence and bleaching. The laser-induced fluorescence can be detected by a photo-optic sensor or a camera. Bleaching is the irreversible destruction of part of the fluorophore. It is assumed that most of the fluorophore is mobile. In our case, we will limit this mobility to standard diffusion. Nevertheless, it is often convenient to assume that a fraction of the fluorophore remains immobile.

In what follows, for each time *t* and each spatial coordinate *r*, we denote the concentration of the unbleached fluorophore by *C*(*r,t*) for the mobile fraction and *C*_im_(*r,t*) for the immobile fraction, respectively. Furthermore, let *I*(*r,t*) be the laser intensity and let *F*(*r,t*) be the corresponding laser-induced fluorescence of the fluorophore.

#### Underlying process model

Let a laser intensity profile *I*(*r,t*) be given. The mobile part of the unbleached fluorophore develops over time according to a reaction-diffusion equation of the form

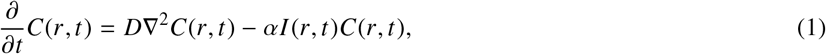

starting with a uniformly constant concentration *C*_0_ at some initial time *T*_0_, i.e. *C*(*r,T*_0_) = *C*_0_ for all *r*. The term *D*∇^2^*C*(*r,t*) on the right hand side of the differential equation describes a standard diffusion process with ∇^2^ being the Laplace operator and *D* being the constant diffusion coefficient of the fluorophore. The term −*αI*(*r,t*)*C*(*r,t*) models a first order bleaching reaction of the fluorophore depending on the rate constant *α* and the laser intensity *I*(*r,t*). The minus sign signifies that bleaching destroys the fluorophore irreversibly.

Similarly, but without the diffusion part, the concentration *C*_im_ of the immobile fraction of the unbleached fluorophore follows the differential equation

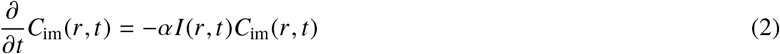

with constant initial condition *C*_im_(*r, T*_0_) = *C*_im,0_ for all *r*.

The fluorescence induced by the laser is then given by

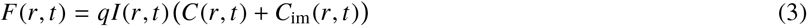

with some constant *q* (which depends on the type of laser and fluorophore).

#### FRAP measurements

The aim of FRAP experiments is to infer knowledge about the diffusion coefficient *D* by inducing a bleaching reaction in a certain region (region of interest, ROI) and the subsequent observation of the time development of the fluorescence inside the ROI. Therefore, three phases of a FRAP experiment can be distinguished: the pre-bleaching phase (*T*_0_ ≤ *t* < −*T*) before bleaching starts at time *t* = −*T*, the bleaching phase (−*T* ≤ *t* < 0) of time duration *T*, and the post-bleaching phase (0 ≤ *t* < ∞) starting at time *t* = 0. Since the bleaching destroys the fluorophore irreversibly inside the ROI, a subsequent observed recovery of the fluorescence can only be due to a transport of unbleached fluorophore from the surroundings of the bleached region via diffusion. The velocity of the recovery allows the computation of estimates of the diffusion coefficient *D*. Here, a model assumption is that the surroundings of the ROI are virtually unbounded. Practically, this means that the probe carrier is large enough and that the ROI is located far from its edges. This guarantees that the amount of fluorophore at the outside of the bleaching region is virtually infinite in each direction, such that the bleached region can fully recover to its initial fluorescence.

#### Laser beam modes

The laser beam operates in two different modes during a FRAP measurement, the bleaching mode and the measurement mode. In both modes, the laser intensity will be based on the same time-independent intensity profile *I*(*r*). In detail:

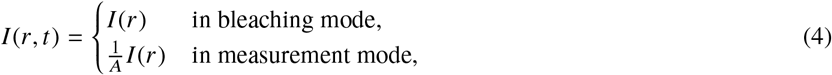

where *A* > 1 is an attenuation factor applied during fluorescence measurement. Practically, the CLSM will be operated in its so-called Flymode, that is, each line is scanned twice: during a forward scan, and only in the bleaching phase, the laser intensity is increased when the laser is located over the ROI, and attenuated when outside, while during the backward scan the laser beam is constantly attenuated in order to perform the fluorescence measurement. With this technique, bleaching and measurement are intertwined during the scanning cycle of each frame, and measurements are obtained in all three phases at equidistant time steps Δ*t* (frame length).

#### Circular intensity profile

We assume that the bleaching region (region of interest, ROI) has a circular two-dimensional shape with radius *w*. Originally, a fixed laser beam was used and the intensity profile was meant to coincide with the laser beam profile itself. In our case, the CLSM offers the possibility to scan a larger ROI with a user-chosen intensity profile. Even though the ROI will be scanned by the CLSM line by line, it will be assumed that the complete region is scanned instantaneously and that each point on that region is scanned with a uniform beam intensity. The circular intensity profile *I*(*r*) and thus the bleaching reaction are symmetric around the origin, such that the coordinates *r* can be given as a (one-dimensional) radial distance from the center of the ROI. The intensity profile is therefore given by

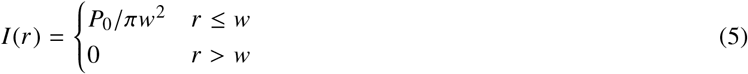

where *P*_0_ is the total laser power over the bleaching region.

#### Bleaching

In order to simplify the computations, further model assumptions have been introduced by (18). The two main assumptions are:

**(A1)** The attenuation factor *A* is high enough such that virtually no bleaching occurs during fluorescent measurement.
**(A2)** The duration time *T* of the bleaching phase is short enough such that virtually no diffusion occurs during that phase.

As a result, always only one part of the diffusion-reaction equation governing the mobile fluorophore concentration is significantly driving the concentration dynamics in each of the FRAP phases. During the bleaching phase,

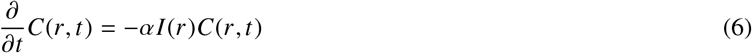

is assumed, leading to the following closed form solution of the concentration at the end of bleaching (*t* = 0):

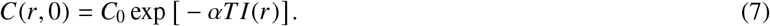

Similarly, the concentration of the immobile fluorophore at time *t* = 0 is given by

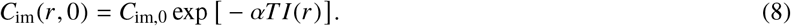

On the other hand, during the pre and post-bleaching phases, despite of the measurements, it is assumed that only diffusion is present for the mobile fluorophore,

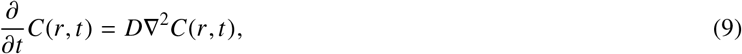

while the immobile fluorophore remains constant at its level *C*_im_(*r*, 0) reached at time *t* = 0.

#### Determination of diffusion coefficient

The total fluorescence *F*(*t*) inside the ROI over time, observed with the attenuated measurement profile 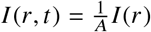, is given by:

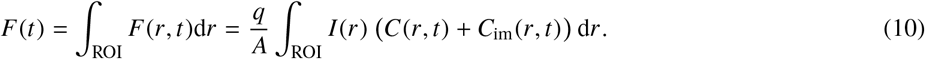

Considering the constant fluorescence values *F*_−_ = *F* (*t*) for *T*_0_ ≤ *t* < *T* (pre-bleaching), *F*0 = *F* (0) (end of bleaching), and *F*_∞_ = lim_*t*→∞_ = *F*(*t*) (asymptotic limit at post-bleaching), *F*(*t*) will be represented as

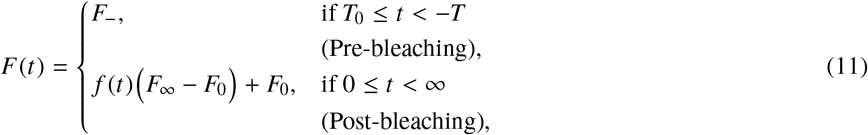

where the fractional fluorescence recovery *f*(*t*), defined as

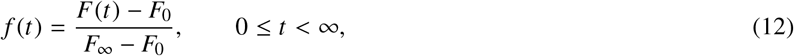

is a function of time with values in the interval [0, 1]. In (18) and (19), it has been shown that *f*(*t*) can be explicitly computed, under the given assumptions and with a circular laser profile, as

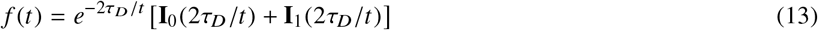

with **I**_0_ and **I**_1_ being the modified Bessel functions and *τ_D_* being the characteristic diffusion time, related to the diffusion constant by

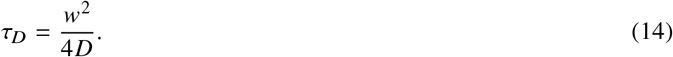

Note that the fluorescence *F*_∞_ after full recovery differs from the fluorescence *F*_−_ before bleaching in the case that part of the fluorophore is immobile. The fraction of total fluorophore in the ROI which is mobile is given by (*F*_∞_ − *F*_0_)/(*F*_−_ − *F*_0_).

#### Measurement data

For each frame *i* of a FRAP measurement cycle, let 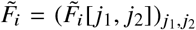 be a matrix representing the intensity values detected during the corresponding laser scan at time *t_i_*. Let *J*_ROI_ be a set of indices (*j*_1_, *j*_2_) representing the pixel coordinates of the ROI, and denote by # *J*_ROI_ its cardinality (number of pixels in the ROI). Then the measurement data yi is given by the mean of the measured intensity values inside the ROI:

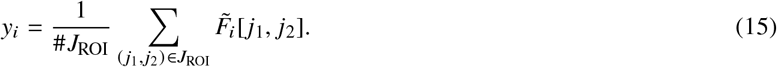

An example of a sequence of fluorescence scans of frames during pre-bleaching and post-bleaching phase on a biofilm probe is shown in Figure 2.

**Figure 2:**
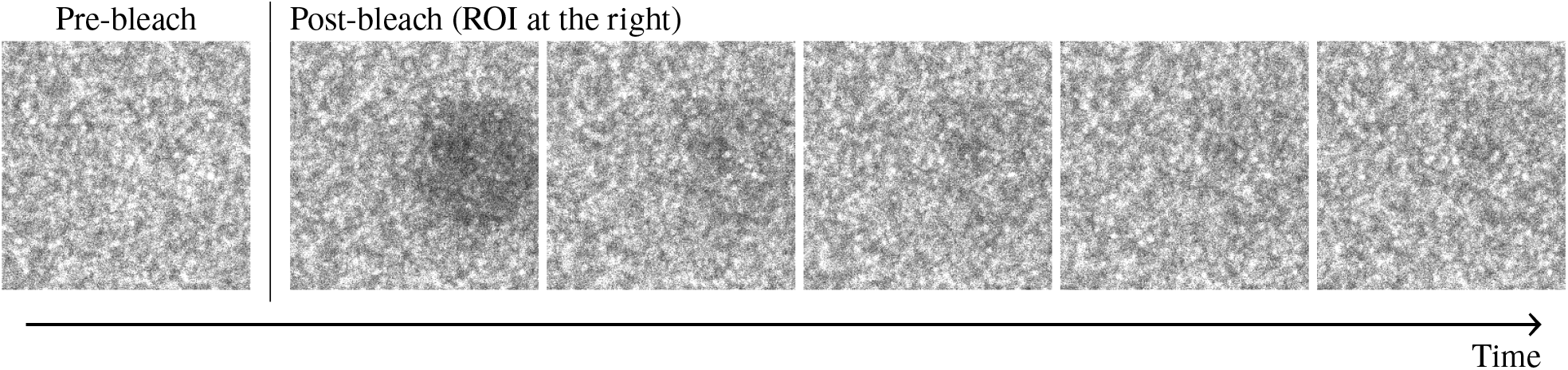
Scanned images during pre-bleaching and post-bleaching phase. The circular ROI is adjacent to the right hand edge of the images and has a radius of 10 *μ*m.

#### Estimation of the diffusion coefficient

Let *i*_0_ be the number of the last frame in the bleaching phase (assumed to be measured at time *t* = 0). Following the model assumptions, it is expected that

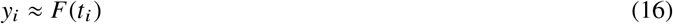

holds for all *i* = *i*_0_,…,*n*. An estimate of the diffusion coefficient *D*can thus be obtained with the following procedure:

1. Normalize the data yi for all *i* = *i*_0_,…,*n* using (estimates of) of *F*_0_ and *F*_∞_ according to

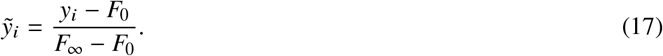
2. Find the parameter *τ_D_* by fitting the curve *f*(*t*) to the normalized data *ỹ_i_*.
3. Set 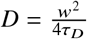.

The parameter search concerning *τ_D_* can be performed in several ways, e.g. by applying a (nonlinear) least squares approach (e.g. steepest descent). Alternatively, the parameters *F*_0_ and *F*_∞_ may be included into the search, in which case one will directly try to fit *F*(*t*) to the unnormalized data *y_i_*.

### FRAP with black-box error model

#### Difficulties with the standard approach

Several difficulties with the standard approach may occur. In order to keep the model equations simple and solvable in closed form, many simplifying model assumptions had to be introduced which do not always apply to the real system. The two main assumptions (A1) and (A2) are often violated. Problems with (A1) may occur if, in order to excite a measurable fluorescence signal, the laser intensity has to be relatively high. This can be the case if for example the overall fluorescence concentration is quite low for some reason. As a result, bleaching during measurements is unavoidable. Problems with (A2) may occur in particular when the diffusion coefficient is high. In fact, the higher the diffusion coefficient, the faster is the fluorescence recovery. In order to compensate for fast recoveries, the time interval Δ*t* between two measurement frames should be decreased together with an eventual increase in the resolution of the ROI. Both actions may be inhibited by limits in the scanning frequencies of the microscope. At the same time, it might be necessary to increase the depth of bleaching which would require to increase the time duration of the bleaching phase. As a result, fluorescence recovery occurs already during bleaching.

Additionally, unmodelled random processes independent from the recovery process may disturb the fluorophore concentration and thus severely interfere with the parameter estimations. In our experiments, we observed that the fluorescence recovery did not asymptotically reach a constant value *F*_∞_ as expected, but rather showed a constant drift, either increasing or decreasing, obviously independent from the recovery process. This effect presumably stems from the technical and biological system employed. A biofilm is never completely homogenous, a microscope can never be fully shielded against environmental influences. A possible cause for the drift could be the very structure of the biofilm - it can have pores and canals, thus causing some kind of mixed diffusion between the dense biofilm and less dense liquid medium. Furthermore, the flow cell is much bigger than the bleaching spot, thus it could be possible that – despite blocking in- and outlet during measurement – there is still some movement in the liquid. It is unlikely that the dye accumulates in the biofilm, which could lead to self-quenching and consequently to a constant increase in fluorescence after bleaching (since there would be less active dye and thus less quenching). Autofluorescence can also be ruled out, since *L. lactis* is not known to have any autofluorescence. In order to fully elucidate the reason for this phenomenon, further studies are to be carried out.

Furthermore, it is not obvious how to get good estimates of the values *F*_0_ and *F*_∞_. It might be tempting to build means over measurements of the pre-bleaching value *F*_−_ as an estimate for *F*_∞_, but if part of the fluorophore is immobile and cannot recover from bleaching, then *F*_∞_ < *F*_−_, and measurements of *F*_−_ cannot be used to predict *F*_∞_. On the other hand, F0 is the fluorescence at the exact time *t* = 0, at the very beginning of the recovery. Shortly after this timepoint, the recovery curve *f*(*t*) is very steep: rapid changes in fluorophore concentration occur because the diffusion rate is very high due to high concentration differences of unbleached fluorophore between the ROI and its surroundings. So, small deviations from the exact point in time of the intensity measurement scheduled for time *t* = 0 may lead to large deviations from the true value of *F*_0_. Additionally, due to the physical process of scanning, there is always some uncertainty concerning the exact times of bleaching and measurements.

In order to cope with these difficulties, we propose to change the traditional approach in two ways: First, we extend the original model by adding an explicit error model to the measurement equation with some new unknown parameters, and secondly, we apply a Metropolis-Hastings algorithm in order to estimate the complete parameter vector.

#### Our extended model

We supplement the FRAP model by adding a simple black-box model which captures several random disturbances in the measurements. Based on our observations, we capture the following phenomena:

- In order to capture effects related to the bleaching of the fluorophore during the fluorescence measurements with attenuated laser intensity, we add an exponential decay term *b*_0_ exp(−*b*_1_*t*) where *b*_0_ is the value of the exponential at time *t* = 0, and *b*_1_ is the rate of exponential decay. This kind of bleaching is most prominently visible at the very beginning of the measurements (i.e. shortly after time *t* = −*T* in the pre-bleaching phase).
- In order to capture the observed drifting of the fluorescence signal, we add a linear drift term *a*_1_*t* with *a*_1_ being the slope.
- We also explicitly add a Gaussian noise term *ση_i_* for each measurement frame *i* where each *η_i_* is independently and standard normally distributed, and where *σ* is the spread (i.e. standard deviation) of the stochastic measurement noise.

Accordingly, the mathematical model for the measurement *y_i_* at time *t_i_* is given by

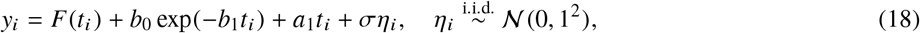

for *i* = i,…,*n*, with new unknown parameters *b*_0_, *b*_1_, *a*_1_ and *σ*, and with *F*(*t*) as defined in Eqs. 11 and 13. It should be noted that we did not include an additional constant offset term in the equation because such a constant can be subsumed under the parameters *F*_−_, *F*_0_ and *F*_∞_ which renders it redundant.

Furthermore, in order to capture the uncertainty concerning the true onset of the post-bleaching phase, the times *t_i_* will be modeled as

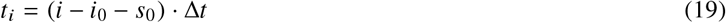

where *i* = 1,…,*n* is the frame number, *i*_0_ the frame number of the last bleaching frame, Δ*t* is the time length of one frame, and *s*_0_ ∈ [0, 1] is a relative time shift which captures the uncertainty in the beginning of the onset of fluorescence recovery.

A prototypical sketch of FRAP measurement values predicted by the original FRAP model, the new error model, and the combined model is shown in Figure 3.

**Figure 3:**
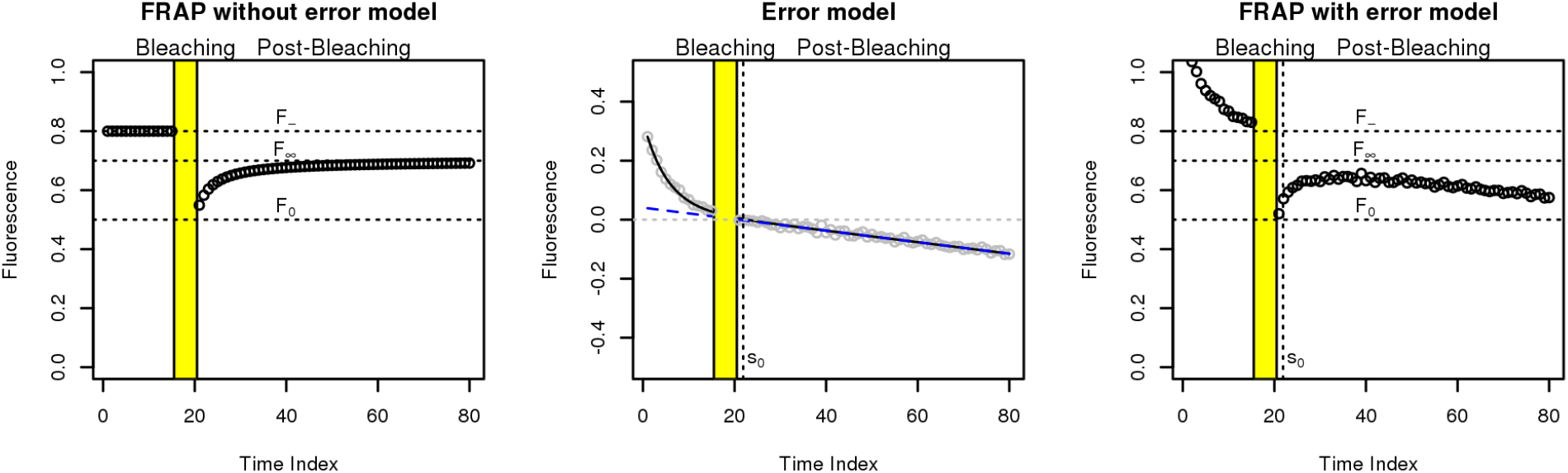
FRAP model (left), error model (middle), and FRAP model with error model combined (right). The error model in the middle panel consists of four parts: 1) a linear drift (parameter *a*_1_, here negative, designated by a dashed line with negative slope), 2) an exponentially decreasing curve which asymptotically approaches the drift (parameters *b*_0_ and *b*_1_, designated by the solid black curve), 3) additive Gaussian noise (parameter *σ*^2^, grey circles), and 4) a time shift (parameter *s*_0_, dotted vertical line). The combined model (right panel) is this error model added to the original FRAP model (left panel). The resulting simulated data look very much like a real FRAP measurement (compare to Figures 6 and 7).

In the following, we will gather all measurements *y_i_* and all unknown model parameters into vectors *y* and *θ*, respectively, i.e.

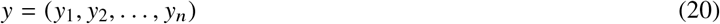

and

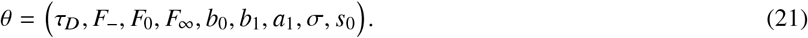

If convenient, wewill denotethesinglecomponents of *θ* by *θ*_1_, *θ*_2_,…,*θ_N_* (where *N* = 9in the extended model).

#### Estimation of diffusion coefficient *D*

In order to estimate the diffusion coefficient *D*, it is necessary to estimate all unknown parameters:

1. Estimate the complete parameter vector *θ* based on the measurements *y*.
2. Set 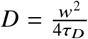

We adopt a Bayesian approach and apply a Metropolis-Hastings algorithm in order to perform the estimation of the unknown parameter vector *θ*. Details of this approach will be given in the following sections.

#### Bayesian approach

The Bayesian model is given by a joint probability density of observations and parameters, *p*(*y, θ*), which factorizes into a product of likelihood *p*(*y*|*θ*) (measurements *y* conditioned on parameters *θ*) and prior density *p*(*θ*):

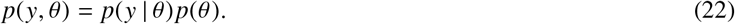

The likelihood is directly given by the model equation Eq. 18 as

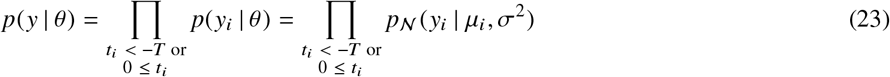

where 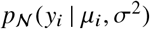 denotes the univariate Gaussian density with mean *μ_i_* and variance *σ*^2^. Here,

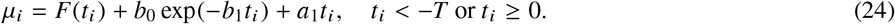

Note that here, in the extended model, the *t_i_*’s are unknown values, too, because they depend on the value of the unknown parameter *s*_0_, according to

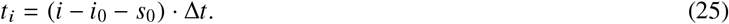

For *i* = *i*_0_, we get *t*_0_ = −*s*_0_Δ*t*, which is practically always strictly less than 0 because the random variable *s*_0_ is practically always strictly greater than 0. For this reason, the measurements corresponding to the last bleaching frame *i*_0_ will not be included into the computation of the likelihood (the same is generally true for all other frames of the bleaching phase: the corresponding measurements will never influence the likelihood).

On the other hand, the prior *p*(*θ*) still needs to be chosen. We decided for an independent uninformative prior, i.e.

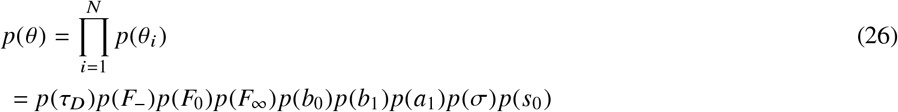

with

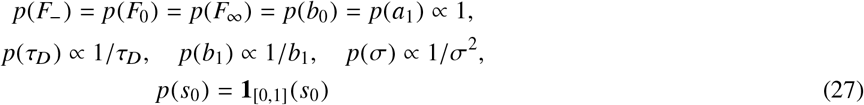

where **1**_[0,1]_ istheindicatorfunction oftheinterval [0, 1].

The target of the estimation procedure is the posterior density *p*(*θ*|*y*), given by Bayes’ theorem as

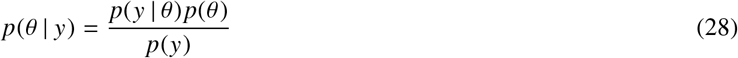

where the marginal data likelihood *p*(*y*) is given by

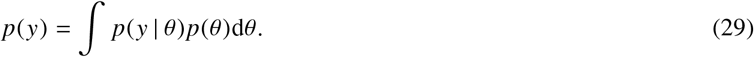

Since the posterior cannot be expressed in a closed form expression, its evaluation must be based on the algorithmic computation of approximations. We applied the Metropolis-Hastings (MH) algorithm, which, given some quite arbitrary proposal distributions (given through probability densities *q_j_* for each parameter *θ_j_, j* = 1,…, *N*), produces a sequence of random samples *θ*^(1)^, *θ*^(2)^, *θ*^(3)^,… of parameter vectors whose distribution approaches the target distribution given by the posterior density *p*(*θ*|*y*).

#### Metropolis-Hastings (MH) algorithm

The Metropolis-Hastings algorithm (30), adapted to our needs, is written in pseudo-code as follows:

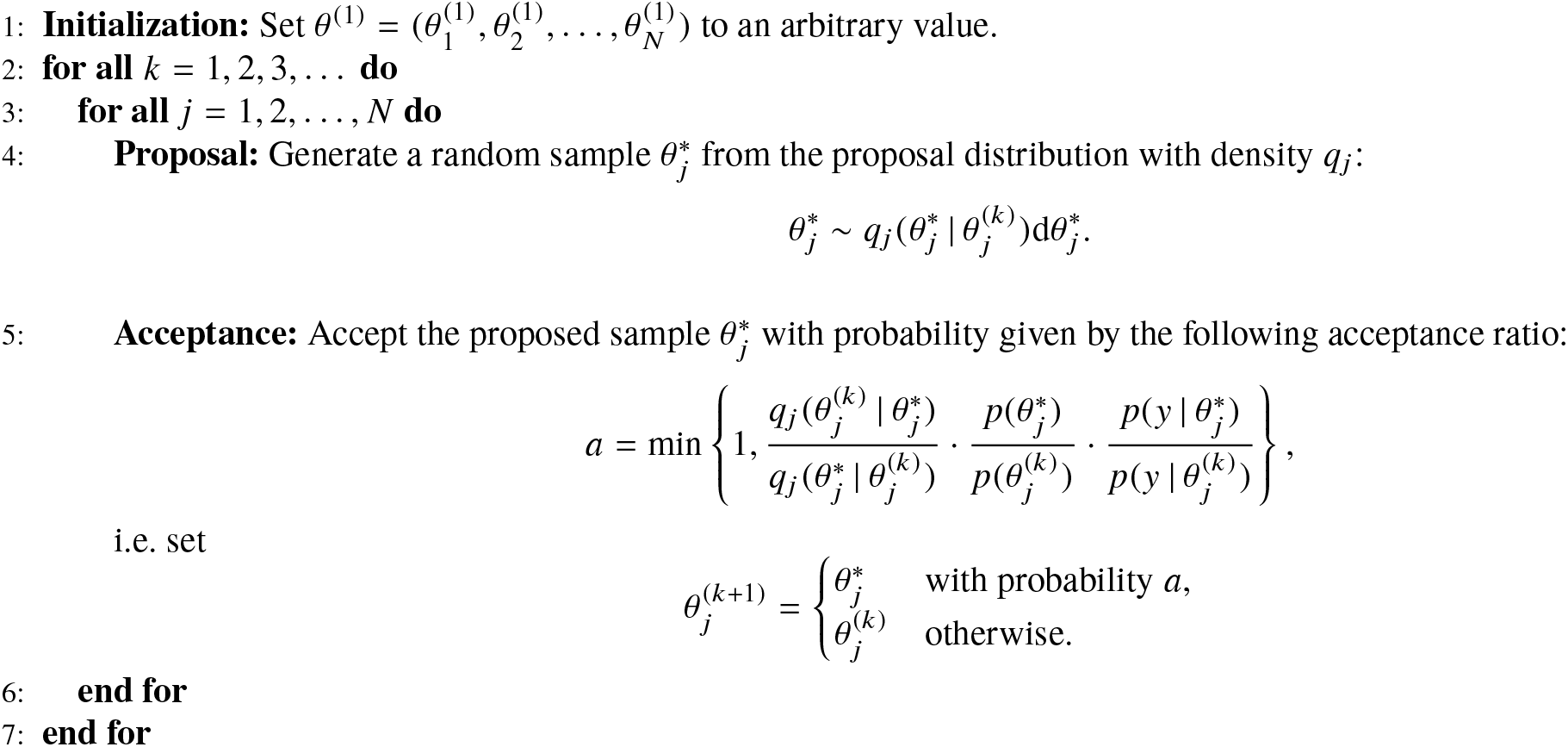

We have implemented the Metropolis-Hastings algorithm from the scratch in the language R (31).

#### Initial values and proposal distributions

The initial parameter vector *θ*^(1)^ has been constructed from crude estimates of the single parameters, i.e. observation means for *F*_0_, *F*_∞_ and *F*_−_, and fixed values for the remaining variables. One loop of our MH algorithm consists of the sequential updating of each of the variables in turn. We used geometric random walk proposals for the positive variables *τ_D_, b*_1_ and *σ*^2^, and (linear) random walk for the remaining variables. The relative time shift *s*_0_ follows a random walk restricted to the interval [0, 1] with reflections at the boundaries. The variances of all these distributions had been fixed after a series of preliminary test runs of the MH algorithm. The exact initial parameter values and proposal distributions can be found in Table 1.

**Table 1:** Initial values and proposal distributions of parameters *θ_j_*. Note that initial *τ_D_* = Δ*T* has been chosen to be the frame length Δ*t* = 0.195[*s*]. The proposal distribution of 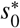 is a normal distribution restricted to [0, 1] with reflection at the boundaries, denoted here by 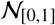. Further, initial 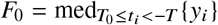 is the median of the total observed fluorescence in the ROI, taken over all bleaching frames.

| $\theta_j$ | Initial value $\theta_j^1$ | Distribution of proposal $\theta_j^*$ |
| --- | --- | --- |
| $\tau_D$ | $\Delta t$ | $\log \mathcal{N}(\log(\tau_D), 0.1^2)$, |
| $F_0$ | $\min_{-T \leq t_i < 0} \{y_i\}$ | $\mathcal{N}(F_0, 0.01^2)$, |
| $F_\infty$ | $\text{med}_{T_0 \leq t_i < -T} \{y_i\}$ | $\mathcal{N}(F_\infty, 0.01^2)$, |
| $F_-$ | $F_\infty$ | $\mathcal{N}(F_-, 0.001^2)$, |
| $a_1$ | 0 | $\mathcal{N}(a_1, 0.001^2)$, |
| $b_0$ | 0 | $\mathcal{N}(b_0, 0.01^2)$, |
| $b_1$ | 1 | $\log \mathcal{N}(\log(b_1), 0.1^2)$, |
| $\sigma^2$ | $0.005^2$ | $\log \mathcal{N}(\log(\sigma^2), 0.1^2)$, |
| $s_0$ | 0.5 | $\mathcal{N}_{[0,1]}(s_0, 0.01^2)$, |

Note that with these choices of proposal distributions, their corresponding ratio terms in the acceptance ratio formula are simply given by

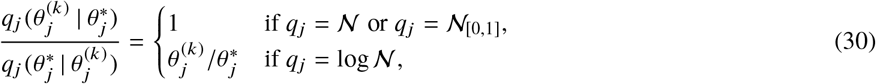

and together with our previous choice of improper prior densities, both ratios will cancel out each other in the computation of the acceptance ratio *a* during each step of the MH algorithm. Thus, the acceptance ratio simplifies to a likelihood ratio:

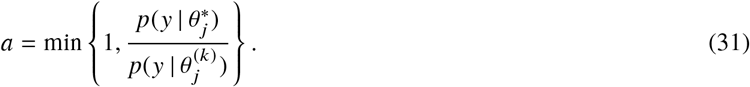

We generally discard the first half of the produced samples (burn-in phase).

## RESULTS AND DISCUSSION

Single FRAP measurements were performed for

- 2 fluorophores: Uranin and FD4,
- 3 positions: pos001, pos002, pos003 (see Figure 1, positions are numbered from left to right following the flow direction),
- 7 times (age of biofilm): at 5h, 7h, 8h, 9h, 10h, 11h, 12h, and
- 2 depths: inside the biofilm, and at the top of it in the medium.

Each of these 84 measurements results in a set of 80 frames (15 pre-bleach, 5 bleach, 60 post-bleach) in the format of TIFF images, which have been directly used to the compute the total fluorescence in the ROI. With these data, Metropolis-Hastings analyses have been performed on two different models, i.e. on a FRAP model 1) with error model and 2) without error model, i.e. in this case, we fixed the parameters *a*_1_, *b*_0_, *b*_1_, *σ*^2^ and *s*_0_ to their initial values given in Table 1 (this specifically means *a*_1_ = *b*_0_ = 0 such that linear drift and exponential decrease are neglected). In this way, the benefits of the error model can be seen by comparison of the two models.

For each single FRAP measurement, we run the Metropolis-Hastings algorithm until 100 000 samples of parameter vectors have been produced. We discarded the first half of the samples (i.e. 50 000) as burn-in, despite the fact that convergence was usually reached already much earlier after a few thousand steps. Accordingly, in each case, all statistical analyses were based solelyon the last 50000 samples.

Since the radius of the ROI is *w* = 10 *μ*m, the diffusion coefficient *D* can be computed from the sampled characteristic times *τ_D_* by

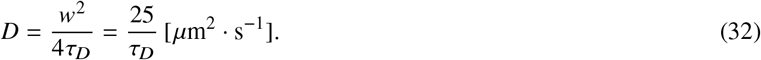

### Estimation results: Overview

In Figures 4 and 5 the results of all 184 MH runs are shown. Each result is represented by a single box-and-whisker diagram. A comparison of the estimation results obtained with error model (left hand sides in both figures) and without error model (corresponding right hand sides) reveals that the additional flexibility present in the error model has an overall positive effect on the results. This is most obvious when the measurements inside the biofilm (dark grey boxes) are considered. On the one hand, the estimations with error model are relatively stable for both uranin and FD4, in the sense that 1) the variation inside each measurement is moderate and nearly equal over all times (5h to 12h), indicated by the nearly equal sizes of the boxes, and 2) the variation of the medians (bars inside the boxes) between different measurements is moderate. On the other hand, the estimations without error model show different aberrations for uranin and for FD4. In the uranin case, the variance within one measurement is lower (i.e. smaller boxes), while the variance between different times is much higher. On the contrary, in the FD4 case, the variance within one measurement is often extremely high (very large boxes), while the median of many measurements (made inside the biofilm) is often higher than the median of the corresponding measurement in the medium. This is against what one would expect because the diffusion coefficient inside the biofilm should be lower than in the medium.

**Figure 4:**
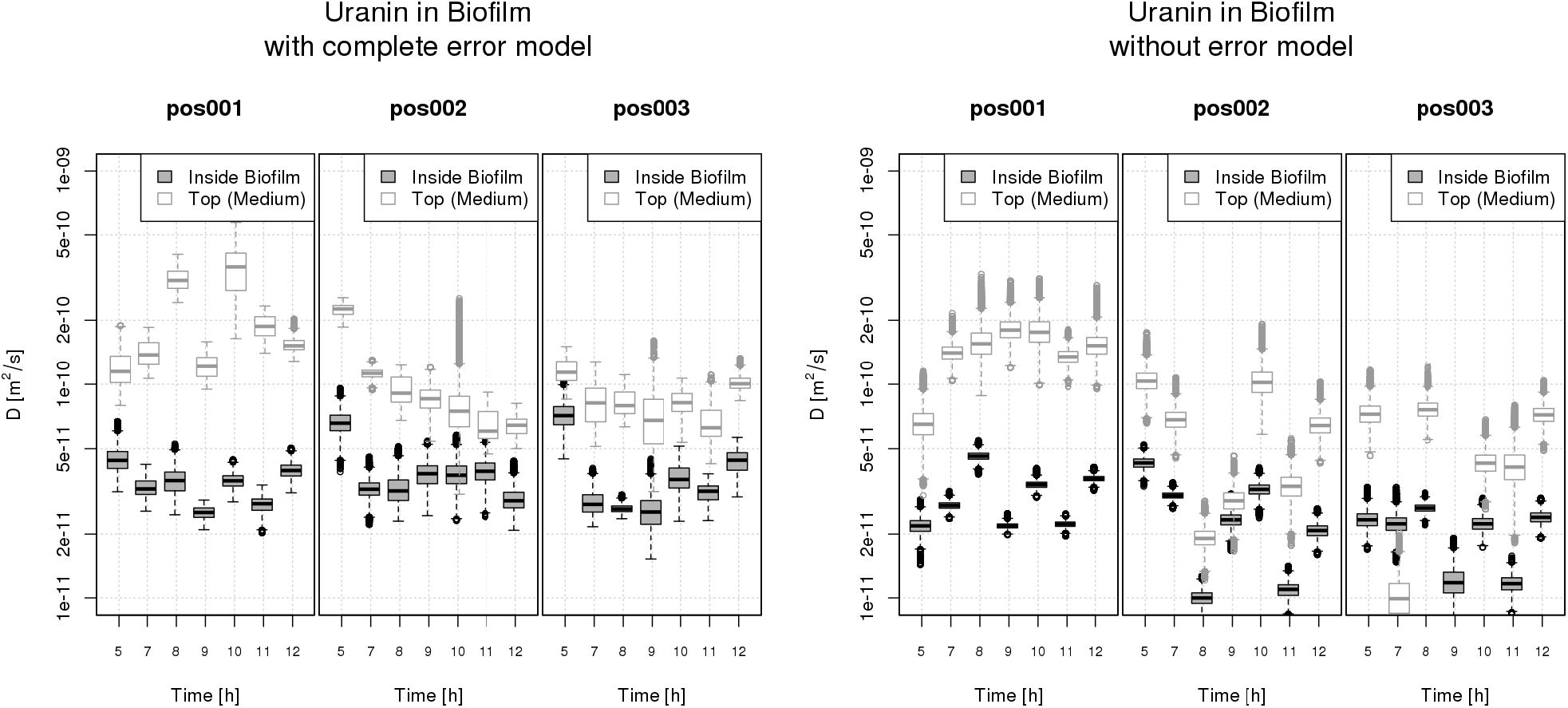
Analysis of FRAP data of uranin diffusion in a biofilm: **with** error model (left) and **without** error model (right). For each FRAP measurement, corresponding MH samples (after burn-in phase) of the diffusion coefficient D are represented by box-and-whisker diagrams. In each single box-and-whisker diagram, the box itself displays the samples between the first and third quartiles (interquartile range, IQR), while the inner band depicts the median. The whiskers extend to the most extreme data point which is no more than 1.5 times the IRQ from the box. The remaining data points (outliers) are plotted as dots.

**Figure 5:**
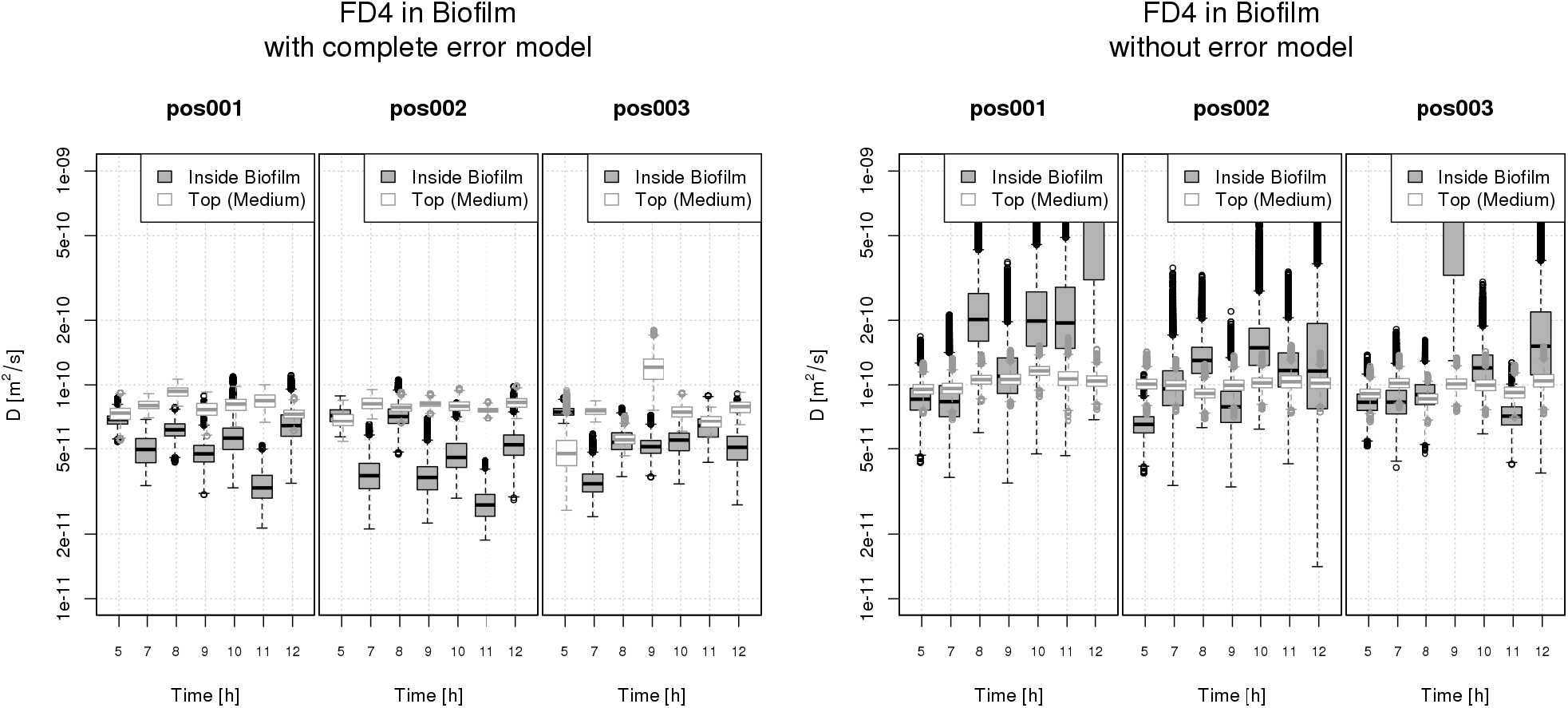
Analysis of FRAP data of FD4 diffusion in a biofilm: **with** error model (left) and **without** error model (right). Description of the box-and-whisker diagrams see Figure 4.

### Two typical examples

We pick out two single FRAP measurements as typical examples.

Our first example is the FD4 measurement at 10h, position 001, measured in the medium (Figure 6). The data show a rapid exponential decrease in the pre-bleaching phase, and a slight but significant negative drift in the post-bleaching phase. Since negative drifts cannot be compensated by adjusting the parameters of the FRAP model, the predictions obtained without the error model do not fit well to the measured data. Nevertheless, despite the bad fit, the estimates of the diffusion coefficients are comparable to the ones estimated with the extended model. On the other hand, the estimates obtained with the extended model provide a much better fit to the data, which shows that the extended model can much better represent the measurements. This in turn heightens the confidence in the validity of the estimated parameters.

**Figure 6:**
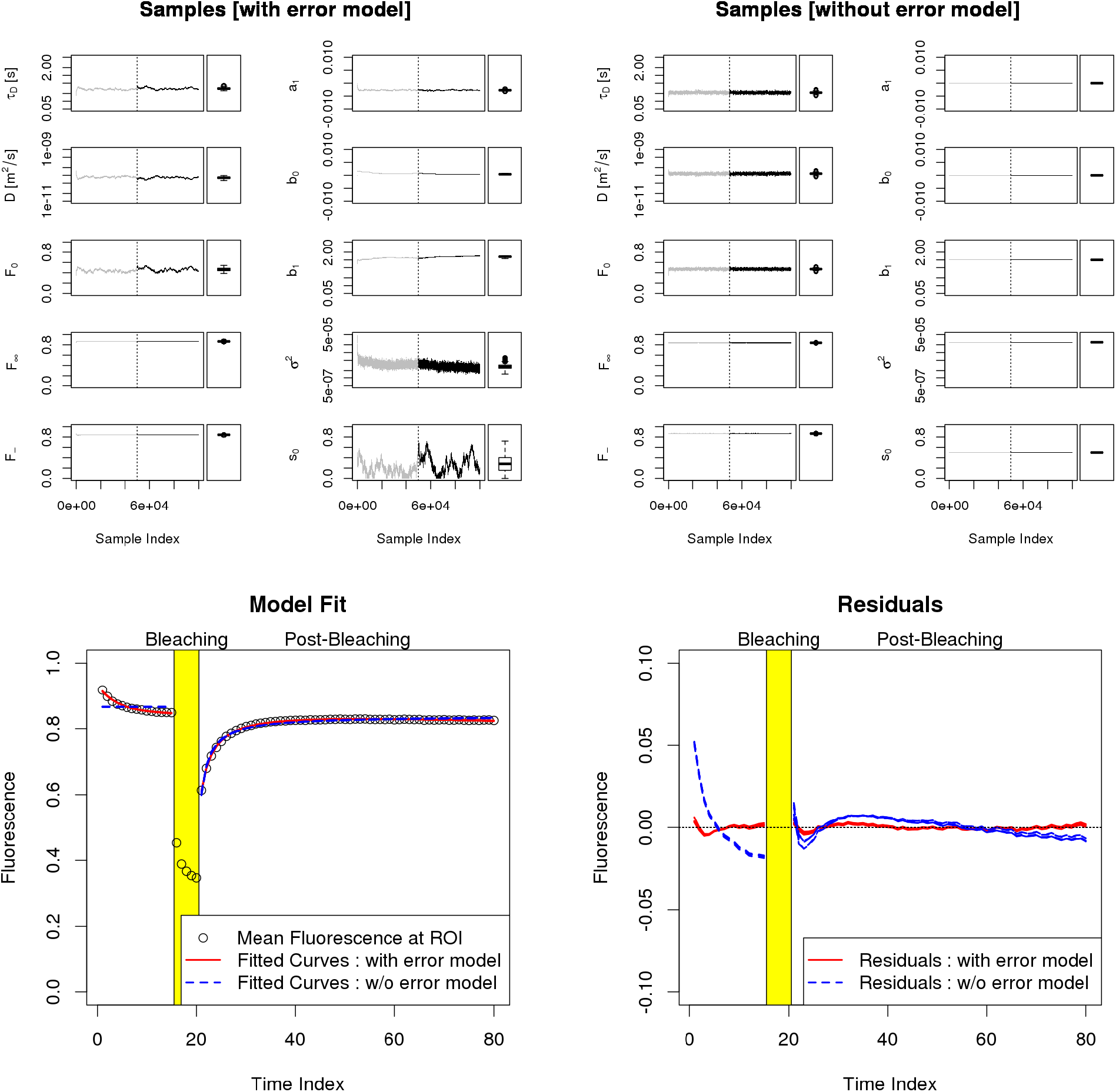
Example 1: FD4 at 10h, position 001, measured in the medium. **Upper row:** Samples produced during MH runs with error model (left) and without error model (right). Shown is the development of the samples over the MH run time, separately for each parameter, and a box-and-whisker diagram (see Figure 4) representing the samples after the burn-in phase. For *τ_D_, D, b*_1_ and *σ*^2^ a log-scale has been used on the y-axis. The end of the burn-in phase has been marked with a vertical line. Samples of burn-in phase are in grey. **Lower row:** Model fit (left) and residuals (right). Shown are the data (black circles), and five model prediction curves with parameters randomly selected from the MH estimations after burn-in, fitted with and without error model, respectively.

The second example is taken from the uranin measurement at 8h, position 002, measured inside the biofilm (Figure 7). Here, the Gaussian noise is higher than in the FD4 measurement of the previous example. The drift in the post-bleaching phase is now positive. Positive drifts can be compensated even in the case when no error model is used, but in this case, the estimated parameters are forced to deviate from their appropriate values in order to obtain this compensation. This leads to an overall good fit of the predicted curve, while the estimated diffusion coefficient is heavily under-estimated. An equally good fit is obtained as well in the case with error model, but here the additional parameters of the extended model capture the compensation. As a result, the estimation of the diffusion coefficient is quite in line with what one would expect. Thus, in this example, too, the confidence in the results obtained with the extended model is much higher.

**Figure 7:**
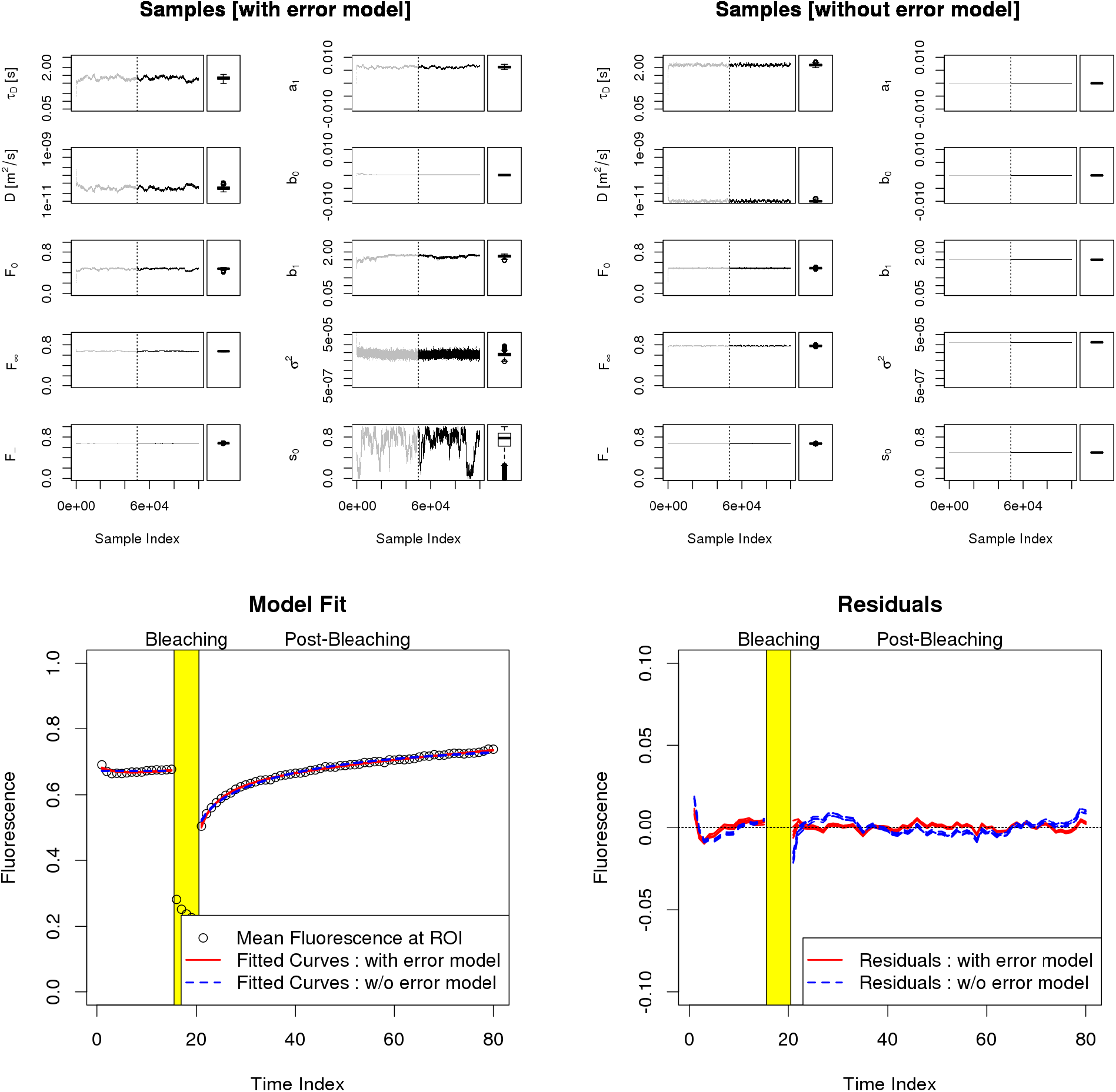
Example 2: Uranin at 8h, position 002, measured inside the biofilm. For further description see Figure 6.

## CONCLUSION

We have extended the standard FRAP model by a black-box part covering some commonly experienced artifacts in the FRAP measurement data, and we have estimated all unknown model parameters (inclusively the diffusion coefficient) through application of a suitably adapted Metropolis-Hastings algorithm. The estimations based on this extended model showed to be much more stable in comparison to estimations obtained without the additional error model. This has been shown on experiments with two different fluorophores in a biofilm. The methodology is not restricted to the application on biofilms and might be easily adapted to other FRAP measurements on other materials. Nevertheless, the proposed model does not capture all disturbances or violations of model assumptions. If one wants to weaken these assumptions, one finally has to abandon the simplified model leading to the basic equation Eq. 13 and has to work with the reaction-diffusion equation Eq. 1 directly. Accordingly, the application of sequential parameter estimation methods might be advantageous or even necessary.

## AUTHOR CONTRIBUTIONS

A.W. and R.U. designed the research. J.C. carried out the experiments and measurements. J.H. analyzed the data. J.H. and J.C. wrote the article.

## ACKNOWLEDGMENTS

This work was funded by the Deutsche Forschungsgemeinschaft (DFG, German Research Foundation) under grants LA 1454/6-1 und UL 170/14-1.

